# Deciphering the landscape of phosphorylated HLA-II ligands

**DOI:** 10.1101/2021.06.29.450288

**Authors:** Marthe Solleder, Julien Racle, Philippe Guillaume, George Coukos, Michal Bassani-Sternberg, David Gfeller

## Abstract

CD4+ T-cell activation in infectious diseases and cancer is governed by the recognition of peptides presented on class II human leukocyte antigen (HLA-II) molecules. Therefore, HLA-II ligands represent promising targets for vaccine design and personalized cancer immunotherapy. Much work has been done to identify and predict unmodified peptides presented on HLA-II molecules. However, little is known about the presentation of phosphorylated HLA-II ligands. Here, we analyzed Mass Spectrometry HLA-II peptidomics data and identified 1,113 unique phosphorylated HLA-II ligands. This enabled us to precisely define phosphorylated binding motifs for more than 30 common HLA-II alleles and to explore various molecular properties of phosphorylated peptides. Our data were further used to develop the first predictor of phosphorylated peptide presentation on HLA-II molecules.

## Introduction

CD4+ T cells play a central role in adaptive immune responses against infections and cancer through the recognition of peptides coming from pathogens or from proteins specifically expressed in cancer cells. The latter include peptides originating from cancer specific genetic or proteomic alterations, often referred to as neo-antigens. Antigen presentation to CD4+ T cells is mediated by class II human leukocyte antigen (HLA-II) molecules, which are expressed on the surface of professional antigen-presenting cells (APCs) such as dendritic cells or B lymphocytes. HLA-II molecules form heterodimers and are encoded by three pairs of genes (HLA-DRA/B, -DPA/B, -DQA/B). Except for HLA-DRA, these genes are highly polymorphic and thousands of alleles have been discovered in humans. HLA-II molecules bind mostly peptides of 12 to 20 amino acids with a 9-mer peptide binding core and flanking regions extending on both sides (see Figure 1A) (Chicz et al., 1992; Neefjes et al., 2011). For most alleles, the binding specificity is driven by the primary anchor residues at P1 and P9 and the secondary anchor residues at P4 and P6 of the peptide core, although some variability has been observed in anchor residues across different HLA-II alleles (Abelin et al., 2019; Racle et al., 2019). HLA-II ligands can originate from both exogenous and intracellular proteins processed by endocytic pathways (Roche and Furuta, 2015), and include both unmodified peptides as well as peptides with post-translational modifications (PTMs) (Lim et al., 2021; Malaker et al., 2017). Recently, HLA-II ligands have been shown to play an important role in the response to personalized cancer vaccines (Alspach et al., 2019; Graciotti et al., 2020; Kranz et al., 2016; Kreiter et al., 2015; Sahin et al., 2017). HLA-II ligands can be identified either by Mass Spectrometry (MS), although such experiments are technically challenging (Caron et al., 2017), or using prediction methods followed by experimental validation. Several predictors of HLA-II ligands have been developed (e.g., NetMHCIIpan (Reynisson et al., 2020a), MixMHC2pred (Racle et al., 2019), or MARIA (Chen et al., 2019)) and can contribute to reduce cost and efforts to identify novel HLA-II ligands, including class II neo-antigens. However, none of the existing predictors specifically integrate PTMs.

**Figure 1:**
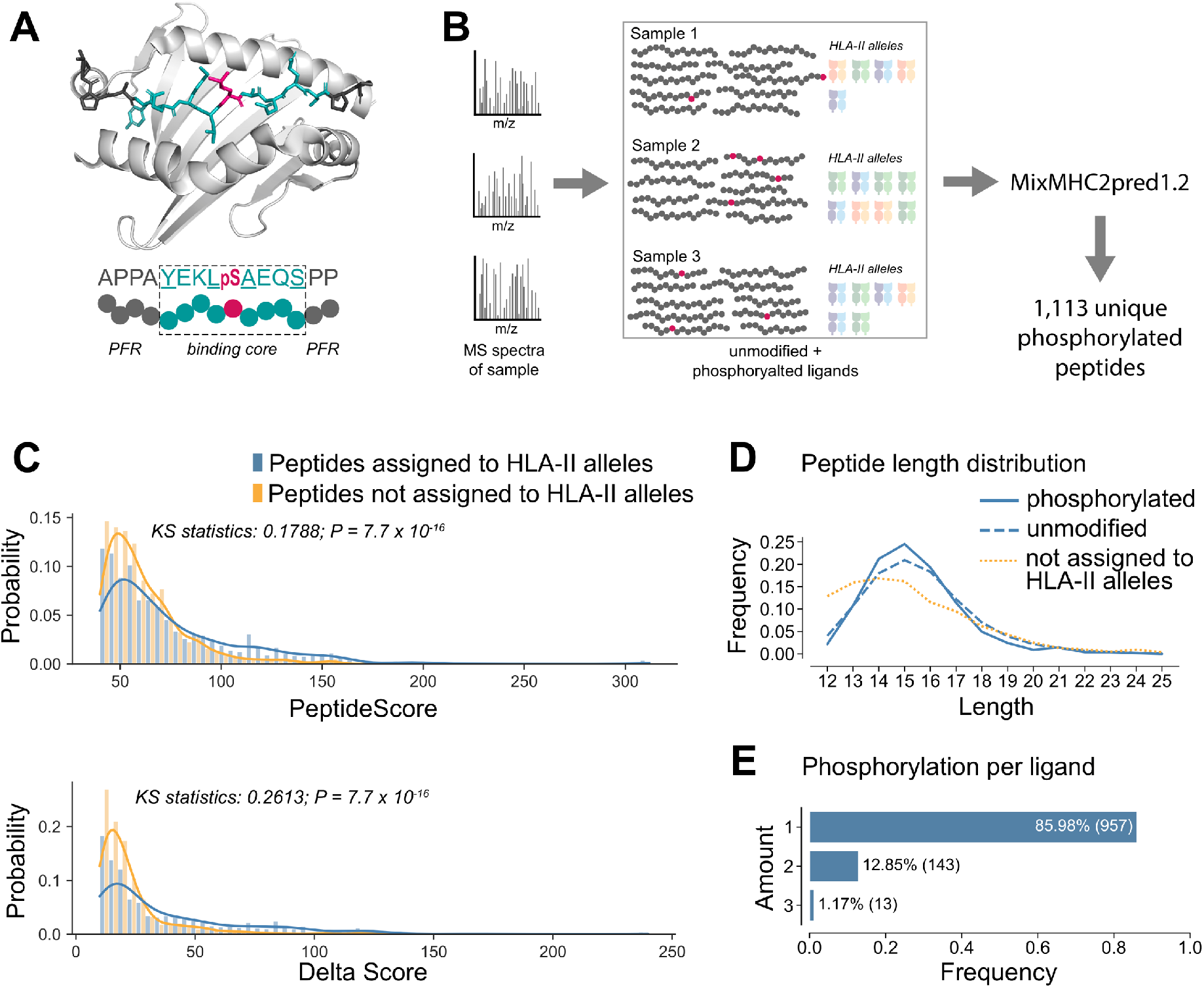
MS-based HLA-II peptidomics identifies multiple phosphorylated HLA-II ligands. **(A)** Representative crystal structure of HLA-DRB1*01:01 molecule in complex with a phosphorylated peptide (PDB identification code 3L6F (Li et al., 2010)). The binding core of the peptide is shown in turquoise, the peptide flanking regions (PFR) in dark grey, the phosphorylated residue in pink, and the HLA-DR in light grey. Anchor positions P1, P4, P6, and P9 are underlined in the peptide sequence and point towards the HLA-II binding site. **(B)** HLA-II peptidomics MS spectra were analyzed for each sample separately to identify HLA-II ligands, including phosphorylated peptides. Phosphorylated peptides were processed for each sample by applying the HLA-II ligand predictor MixMHC2pred. **(C)** Distribution of Andromeda search engine peptide spectrum match scores (*Peptide Score*) (top) and score differences to the second-best peptide spectrum match (*Delta Score*) (bottom) of phosphorylated peptides assigned to HLA-II alleles (blue) and phosphorylated peptides not assigned to any allele (orange). **(D)** Comparison of length distribution of phosphorylated and unmodified HLA-II ligands as well as predicted phosphorylated peptides not assigned to HLA-II alleles. **(E)** Amount of detected phosphorylated residues per phosphorylated HLA-II ligand in the HLA-II phosphopeptidome.

PTMs of proteins are essential regulators in many biological processes (Graves and Krebs, 1999; Hunter, 2009; Wang et al., 2014). PTMs like phosphorylation were shown to be deregulated in cancer cells, causing aberrant cellular behavior (Krueger and Srivastava, 2006; López-Otín and Hunter, 2010; Martín-Bernabé et al., 2017). Therefore, phosphorylated peptides presented on HLA molecules provide potential targets for the development of immunotherapeutic strategies (Engelhard et al., 2020; Lin et al., 2019; Meyer et al., 2009; Petersen et al., 2009a). While many studies analyzed phosphorylated peptides presented on HLA-I molecules (Alpízar et al., 2017; Andersen et al., 1999; Cobbold et al., 2013; Mohammed et al., 2008; Petersen et al., 2009b; Solleder et al., 2020; Zarling et al., 2006), phosphorylated HLA-II ligands have received much less attention. The first naturally presented phosphorylated HLA-II ligands were identified from an EBV – transformed B-lymphoblastoid and a tumor cell line (Meyer et al., 2009). Shortly after, the first CD4+ T cell recognition of a phosphorylated HLA-II ligand was shown using the melanoma antigen Melan-A/MART-1 (Depontieu et al., 2009). Structural analysis of a phosphorylated peptide bound to HLA-DRB1 showed that the phosphorylated residue can in this case directly interact with the T-cell receptor (Li et al., 2010). While these studies provide evidences for HLA-II presentation of phosphorylated peptides and show potential applications as targets for immunotherapies, further characteristics such as binding motifs of phosphorylated HLA-II ligands on a large allelic coverage remain unknown and no HLA-II ligand predictor is specifically trained on phosphorylated ligands.

In this work, we reprocessed 23 high quality MS HLA-II peptidomics samples including phosphorylation in the spectral searches and identified 1,113 novel phosphorylated HLA-II ligands. Based on this data, we defined phosphorylated binding motifs of HLA-II alleles and identified specific molecular properties of phosphorylated HLA-II ligands. Furthermore, we developed the first HLA-II ligand prediction method specifically trained on phosphorylated peptides.

## Results

### MS-based HLA-II peptidomics identifies multiple phosphorylated HLA-II ligands

To identify a broad spectrum of phosphorylated HLA-II ligands across a wide range of HLA-II alleles, we reanalyzed raw MS HLA-II peptidomics data of 23 polyallelic samples (Racle et al., 2019). 13 of these samples were generated with HLA-DR and pan-HLA-II antibodies and the other 10 only with pan-HLA-II antibodies. We used MaxQuant, allowing for phosphorylation on serine, threonine, and tyrosine as variable modifications (see Methods). To ensure broad coverage, we chose a loose false-discovery rate (FDR) of 5% with restricted peptide identification scores (see Methods). Across all samples, a total of 3,471 phosphorylated peptides (representing 2,800 unique peptides) were detected (Table S1). To determine HLA-II allelic restriction and remove expected contaminants or wrongly identified peptides, we performed predictions for each allele of each sample where the peptide was found using the HLA-II ligand predictor MixMHC2pred v1.2 (Racle et al., 2019). For these predictions, phosphorylated residues were treated as ‘X’ in MixMHC2pred (see Figure 1B and Methods). 1,233 HLA-II – phosphorylated peptide interactions (representing 1,113 unique peptides, see Table S2) passed the %rank threshold of 5%, which is commonly used to determine weak binders (Reynisson et al., 2020b). The other cases may consist of co-eluted contaminants or wrongly identified peptides, as expected in HLA-II peptidomics data (Racle et al., 2019). To support this hypothesis, we compared both the distribution of the scores for peptide spectrum matches from the Andromeda search engine (*Peptide Score*, higher values for higher confidence in peptide identification) and the distribution of the score differences to the second best peptide spectrum match (*Delta Score*, higher values for unambiguous distinction from other peptides) for peptides assigned to HLA-II alleles (blue in Figure 1C) and peptides that did not pass the MixMHC2pred filtering (orange in Figure 1C) (see also Figure S1A). As expected, phosphorylated peptides that could not be assigned to any HLA-II allele showed lower *Peptide Scores* (KS statistics: 0.1788; P=7.7 × 10^−16^) and lower *Delta Scores* (KS statistics: 0.2613; P= 7.7 × 10^−16^) than those that could be assigned to HLA-II alleles. These peptides were therefore excluded from downstream analyses. This represents more than half (58%) of the peptides identified in our different samples. The same filtering applied to random peptides selected from a pool of all known phosphosites of the human proteome (Ullah et al., 2016) resulted in the exclusion of 80.8% of the peptides (see Methods). This demonstrates that our set of phosphorylated peptides is enriched in peptides matching HLA-II motifs. The set of 1,113 unique phosphorylated HLA-II ligands showed a length distribution similar to the one of unmodified HLA-II ligands with a peak around 15-mers, while the phosphorylated peptides not assigned to any allele and therefore excluded from downstream analyses had a length distribution skewed towards shorter peptides (Figure 1D). The majority of phosphorylated HLA-II ligands contained one phosphorylated residue, while 12.85% of the peptides are double phosphorylated and ~1% triple phosphorylated (Figure 1E).

#### Phosphorylated peptides bind to HLA-II molecules with specific motifs

The 1,113 phosphorylated peptides could be assigned to 33 different alleles (Table S2). We then used the predicted binding cores to build binding motifs of phosphorylated HLA-II ligands for each of these alleles. As expected, the binding motifs showed conserved specificity at anchor residues P1, P4, P6, and P9 for most alleles (see Figure 2). We observed similar frequency of phosphorylated peptides for different HLA-II genes, with the exception of the two HLA-DRB3 and the two HLA-DRB4 alleles which had higher fraction of phosphorylated ligands (Figure S1B). This enrichment may be explained by specificity for aspartic acid at P4 in HLA-DRB3 alleles and P7 in HLA-DRB4 alleles, which could more easily accommodate phosphorylated residues compared to hydrophobic anchor residues found in many other alleles (Figure 2).

**Figure 2:**
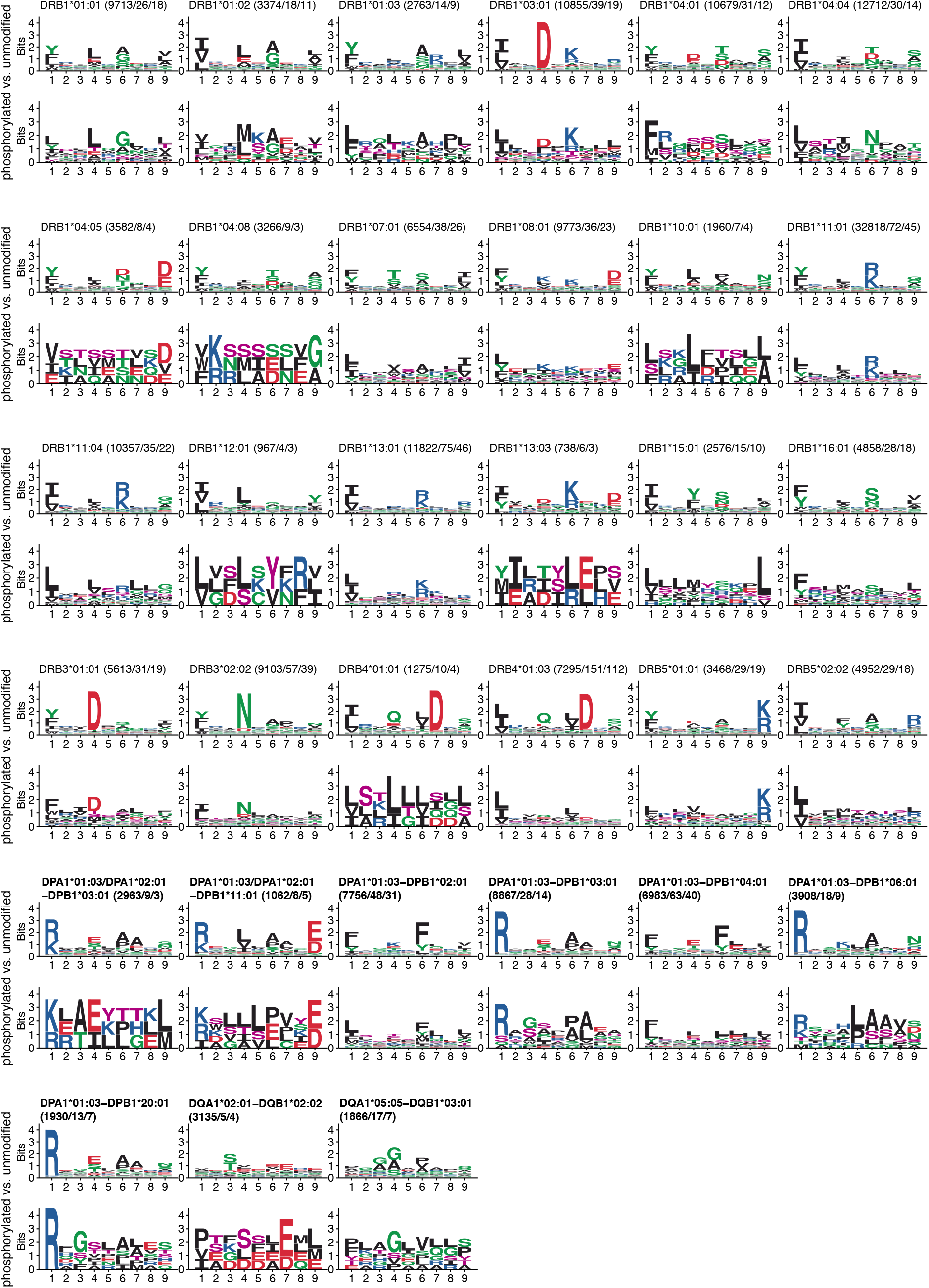
Phosphorylated peptides bind to HLA-II molecules with specific motifs. Motifs of alleles with phosphorylated peptides. For each allele, the HLA-II motif based on unmodified ligands is shown on top, and the motif of phosphorylated HLA-II ligands determined in this work is shown below. Numbers correspond to the number of peptides (unmodified peptides / all phosphorylated peptides / only phosphorylated peptides with the phosphorylated residue in the core). Phosphorylated residues are shown in pink.

#### Phosphorylated residues show positional specificity in HLA-II ligands

To investigate whether there is any preference for phosphorylated residues in the peptide binding core and the peptide flanking regions (PFRs), we compared the fraction of phosphorylated residues with the total fraction of residues in these two regions of the peptide. We could see that phosphorylated residues show a slight enrichment in the PFRs (Figure 3A, P=0.025). We then analyzed phosphorylated residues in PFRs and compared their frequency in the first and last three amino acids of the PFRs at the N- and C-terminus of the phosphorylated peptides. Specific preferences for amino acids in these regions have been attributed to peptide cleavage and processing (Barra et al., 2018; Ciudad et al., 2017; Racle et al., 2019). Phosphorylation sites occurred more frequently at the C-terminus of phosphorylated HLA-II ligands than at the N-terminus (with 61% and 39% of terminal phosphorylation found at C- and at the N-terminus, respectively). We hypothesized that this could be due to the presence of clearer cleavage motifs at the N-terminus, and especially the preference for proline at the second position (Racle et al., 2019). This hypothesis is consistent with the distribution of phosphorylated residues within the N-terminal region where phosphorylation is mostly found at the third position (46.6%) and less at the other two (27.2% and 26.2%, respectively) (Figure 3B). We then looked at the distribution of phosphorylated residues within the 9-mer binding core. We could clearly see less phosphorylated residues at the anchor position P1, which is consistent with the high specificity observed at this main anchor position in unmodified HLA-II ligands, mainly for hydrophobic residues (Figure 3C). The highest frequency of phosphorylated residues is seen at the non-anchor position P5, which shows low specificity in HLA-II binding motifs. Other positions such as at secondary anchor positions (especially P4 and P6) show more variability in the unmodified HLA-II binding motifs, which is also reflected by the presence of phosphorylated residues observed at these positions.

**Figure 3:**
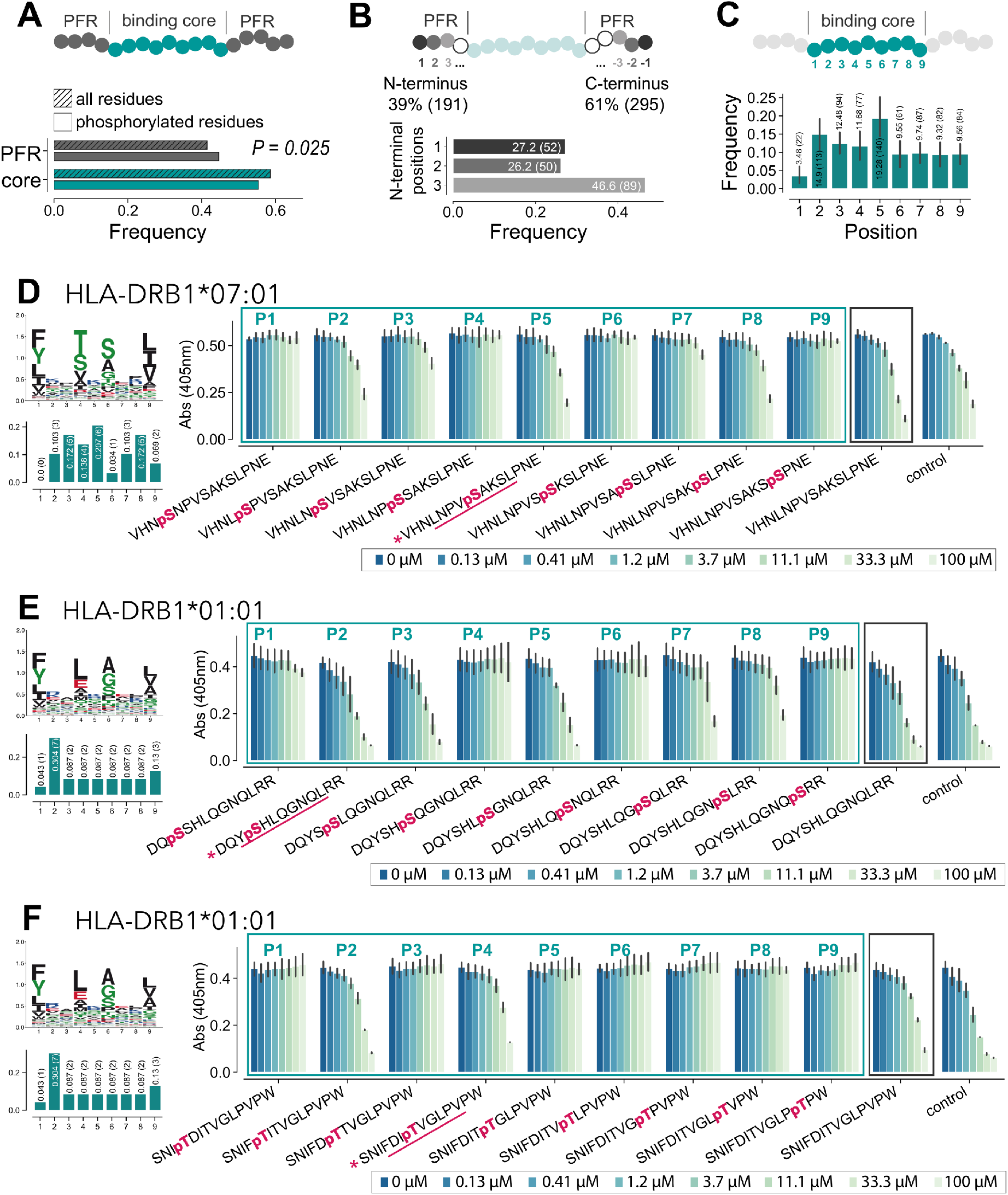
Phosphorylated residues show some positional specificity in HLA-II ligands. **(A)** Distribution of phosphorylated residues and total residues in the binding core vs PFRs of phosphorylated HLA-II ligands. **(B)** Amount of phosphorylated residues found in the first three and last three residues in PFRs (top) and distribution of phosphorylated residues within the first three positions of the N-terminus (bottom) of phosphorylated HLA-II ligands. **(C)** Positional distribution of phosphorylated residues in the binding core of phosphorylated HLA-II ligands. **(D-F)** Competitor binding assays for peptides with a phosphorylated residue at each of the different core positions (turquoise box) and without phosphorylated residue (black box). The peptide initially found by MS is marked by a pink asterisk and the core predicted by MixMHC2pred is underlined. HLA-II motif and positional distribution of phosphorylated residues in the binding core of phosphorylated ligands of the respective allele are shown on the left.

To further investigate the preference for phosphorylated residues at specific positions in the core, we performed competitor binding assays for two different HLA-DR alleles testing different versions of the same peptide containing the phosphorylated residues at all possible positions within the core (see Methods). The two peptides were selected among the set of phosphorylated HLA-II ligands identified by MS with the phosphorylated residue at the non-anchor positions P2 and P5, respectively (Figure 3D-E). The results of the binding assays showed that for both alleles, the peptide that was found in our MS data displayed good binding (see Figure 3D for HLA-DRB1*07:01 with pS at P5 and Figure 3E for HLA-DRB1*01:01 with pS at P2). The unmodified version of the peptide was binding equally well. The presence of the phosphorylated residue at other positions showed inferior binding, especially at positions P1, P4, P6, and P9. These positions could clearly be identified as anchor positions of the alleles (see binding motifs Figure 3D, E left panels). We then selected another peptide for HLA-DRB1*01:01 that was found in our MS data with a phosphorylated residue predicted at the secondary anchor position P4. The low binding with the phosphorylated residue at P1 and P9 could be confirmed. However, for other core positions the results did not reflect the anchor residues and binding of this peptide was detected with a phosphorylated residue at P2 and P4, but not at P5, for instance (Figure 3F). Overall, these observations suggest that the preference for the position of the phosphorylated residue in the middle of the core may be different for different peptides, which could explain the relatively broad distribution in Figure 3C, and the lack of exclusion of P4 and P6 secondary anchor positions.

#### Kinase motifs in HLA-II ligands

To investigate the presence of kinase motifs in the HLA-II phosphopeptidome, we searched for known kinase motifs from the PhosphoMotif Finder of the Human Protein Reference Database (Amanchy et al., 2007) both in phosphorylated and unmodified HLA-II ligands as well as the human phosphoproteome (Sharma et al., 2014) (see Methods). Motifs that show a significant enrichment between phosphorylated and unmodified HLA-II ligands (P ≤ 0.05) are shown in Figure 4A. These include frequent kinase motifs like [K/R]XX[pS/pT] or [pS/pT]XX[D/E]. However, not all common kinase motifs were enriched in our data. For instance, the frequent kinase motif [pS/pT]P, which corresponds to proline-dependent serine/threonine kinases such as MAPK1, was only detected for 6.68% of phosphorylated serine and threonine in our phosphorylated HLA-II ligands. This corresponds roughly to the frequency of proline after serine or threonine in the human proteome (6.78%) or in unmodified HLA-II ligands (5.99%), and stands in contrast to the observed frequency of ~32% at phosphorylated serine and threonine in the human phosphoproteome. To assess what could be the reasons for the lack of enrichment of this kinase motif in the HLA-II phosphopeptidome, we investigated if this may reflect a source protein bias in the HLA-II peptidome (i.e., peptides coming from proteins with such phosphorylation sites would be under-represented in HLA-II ligands, irrespective of the phosphorylation status). To this end, we computed the overlap between the source genes of all HLA-II ligands and the source genes of proteins containing phosphosites with the [pS/pT]P motifs in the human phosphoproteome. This overlap was the expected one (odds ratio: 0.988), suggesting that there is no depletion of proteins containing the [pS/pT]P motifs in the HLA-II peptidome (see Figure S2). This supports the idea that such phosphosites are present in their unphosphorylated form in our set of HLA-II ligands.

**Figure 4:**
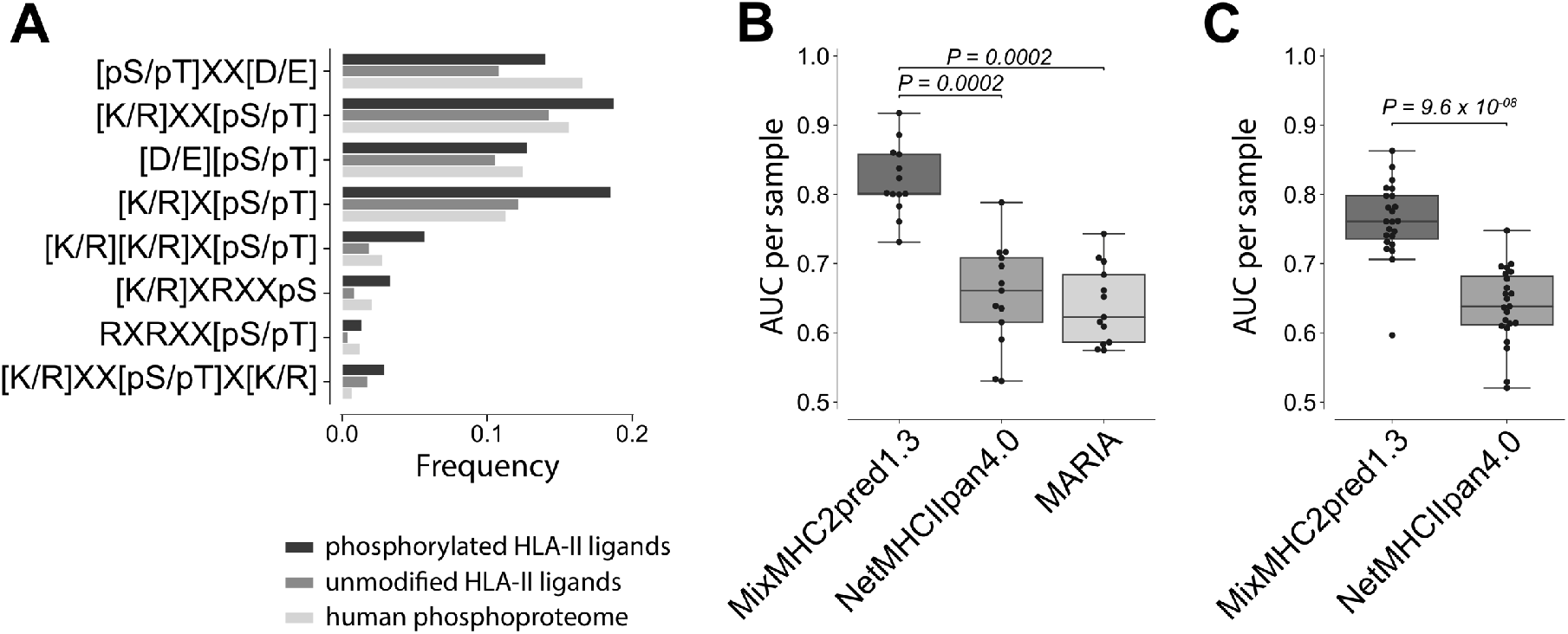
The HLA-II phosphopeptidome improves prediction of phosphorylated HLA-II ligands. **(A)** Frequency of kinase motifs that show significant enrichment between phosphorylated and unmodified HLA-II ligands (P ≤ 0.05) in phosphorylated HLA-II peptidome (1^st^ bar), in unmodified HLA-II ligands (2^nd^ bar) and in the human phosphoproteome (3^rd^ bar). Kinase motifs are sorted according to the frequency in the human phosphoproteome. **(B)** AUC values for the leave-one-sample-out cross-validation for all HLA-DR samples processed in this work (13 in total) for MixMCH2pred1.3, NetMHCIIpred4.0, and MARIA. P-values between the different predictors were calculated using the paired two-sided Wilcoxon signed rank-test. **(C)** AUC values for the leave-one-sample-out cross-validation for all HLA-II samples processed in this work (23 in total) for MixMHC2pred1.3 and NetMHCIIpan4.0.

#### The HLA-II phosphopeptidome improves prediction of phosphorylated HLA-II ligands

We used the HLA-II phosphopeptidome to expand our HLA-II ligand prediction method (MixMHC2pred (Racle et al., 2019)) to phosphorylated peptides. To this end, MixMHC2pred was retrained combining both unmodified and phosphorylated peptides (see Methods). To benchmark its performance, we performed a leave-one-sample-out cross-validation for each sample in our dataset. In each round of the cross-validation one sample was used as test set and the training of MixMHC2pred was done with all phosphorylated peptides that could be assigned to alleles, excluding those that were present in the sample used as test set (see Methods). For each sample, phosphorylated peptides used as negatives were randomly selected from the human phosphoproteome, in 5-fold excess compared to the positives (see Methods). Importantly, the sample used as test set was not filtered with any predictor to prevent potential biases in the comparison between the different predictors. The area under the receiver operating characteristic curve (AUC) was used to assess the performance and to benchmark the new version of MixMHC2pred (v1.3) with the existing tools NetMHCIIpan4.0 (Reynisson et al., 2020a) and MARIA (Chen et al., 2019). In the first benchmark, we restricted to samples measured with HLA-DR specific antibodies, since MARIA predictions are limited to HLA-DR alleles. Both, NetMHCIIpan4.0 and MARIA are not specifically trained for modified residues, thus predictions of phosphorylated peptides with these tools were performed by substituting phosphorylated residues with ‘X’ for NetMHCIIpan4.0 and by using the unmodified counterpart of the phosphorylated residue for MARIA (‘X’ are not supported in MARIA). Figure 4B shows improved predictions with MixMHC2pred. In a second benchmark, we considered data from all samples used in this work and restricted the comparison to NetMHCIIpan4.0. Here again, MixMHC2pred1.3 displayed improved accuracy (Figure 4C).

## Discussion

A better understanding of the repertoire and the properties of HLA-II ligands is promising for the development of personalized cancer immunotherapies such as cancer vaccines (Alspach et al., 2019; Kranz et al., 2016; Kreiter et al., 2015; Sahin et al., 2017). As cancer can cause aberrant PTMs, large datasets of modified HLA-II ligands informing us about the rules for the presentation of such ligands and enabling us to predict them are useful to expand the list of potential targets for cancer immunotherapy. It is also likely that several phosphorylated peptides from pathogens are displayed on HLA-II molecules, although little data is available about them, partly because of the previous lack of predictors for such ligands.

In this work, we performed an in-depth analysis of the HLA-II phosphopeptidome. We could identify binding motifs of phosphorylated HLA-II ligands for more than 30 alleles. These binding motifs showed high similarity with those of unmodified HLA-II ligands at anchor positions, in particular the main anchors at P1 and P9.

Our analysis of the position of phosphorylated residues in HLA-II ligands revealed a preference for phosphorylation in the middle of the core (P5), and low frequency of phosphorylated residues at the anchor position P1 (Figure 3A, C). These results could be confirmed with binding assays and are consistent with the low frequency of phosphorylated residues at anchor positions in HLA-I ligands (Solleder et al., 2020). The presence of phosphorylated residues at secondary anchor positions (mainly P4 and P6) was less expected. However, our binding assays confirmed that specific alleles can accommodate phosphorylated residues at such secondary anchor positions (Figure 3F). The lower frequency of phosphorylated residues at the N-terminus compared to the C-terminus of the HLA-II ligands as well as the depletion of phosphorylation at the first and second positions of the N-terminus (Figure 3B) suggest that the presence of phosphorylated residues at these positions may not be favorable for protein cleavage or transport, although additional work will be needed to confirm this hypothesis.

Our analysis of kinase motifs detected over-representation of only a few known kinase motifs. For instance, the very frequent [pS/pT]P motif was not seen at higher frequency in the phosphorylated HLA-II peptidome compared to the unmodified HLA-II peptidome. We speculate that many of these potential phosphosites are simply not phosphorylated in the pool of ligands available for loading onto HLA-II molecules or that the phosphate groups are removed by phosphatases before or after binding to the HLA-II molecules. The limited enrichment in motifs for intracellular kinases also supports the idea that phosphorylated residues observed among HLA-II ligands come from a more diverse repertoire of kinases compared to the one observed in the HLA-I phosphopeptidome (Solleder et al., 2020). This hypothesis is consistent with the differences between class I and class II antigen presentation pathways and the fact that many HLA-II ligands come from endocytosis of proteins in the extracellular matrix, which may undergo phosphorylation by different sets of kinases compared to intracellular proteins displayed on HLA-I molecules.

To facilitate further studies of phosphorylated HLA-II ligands, we used our data to build a predictor for phosphorylated HLA-II ligands by including the HLA-II phosphopeptidome in the training data of our HLA-II ligand prediction method MixMHC2pred (v1.3). Our results show that this new predictor has higher accuracy compared to other tools (Figure 4B, 4C). The motifs of phosphorylated HLA-II ligands suggest that the binding of phosphorylated peptides is shaped by the binding motif of the HLA-II allele and some positional specificity for the phosphorylated residues (e.g., exclusion of P1), and that this information is accurately captured by MixMHC2pred1.3.

Altogether, our work represents the first in-depth analysis of the repertoire of phosphorylated HLA-II ligands. We anticipate that this resource and the associated computational tools to predict phosphorylated HLA-II ligands in different contexts will facilitate the discovery of potential new targets for CD4+ T-cell recognition in infectious diseases and cancer immunotherapy.

## Methods

### Curation of immunopeptidomics HLA-II MS datasets

The MaxQuant platform (Cox and Mann, 2008) version 1.5.5.1 was employed to search the MS peak lists of 23 samples from (Racle et al., 2019) against a fasta file containing the human proteome (Homo_sapiens_UP000005640_9606, the reviewed part of UniProt, with no isoforms, including 21,026 entries downloaded in March 2017) and a list of 247 frequently observed contaminants. Peptides with a length between 8 and 25 amino acids were allowed. The second peptide identification option in Andromeda was enabled and the enzyme specificity was set as unspecific. An FDR of 5% was required for peptides and no protein false-discovery rate was set. The initial allowed mass deviation of the precursor ion was set to 6 ppm and the maximum fragment mass deviation was set to 20 ppm. Methionine oxidation, N-terminal acetylation and phosphorylation on serine, threonine, and tyrosine were set as variable modifications. The resulting list of msms identifications were further filtered to include phosphorylated peptides with identification score ≥ 40, score difference to the second best peptide spectrum match (*Delta Score*) ≥ 10, and localization probability for phosphorylation of >0.75 as well as peptide lengths restricted to 12 to 25 amino acids (Table S1). To obtain better specificity for unmodified peptides, sample-specific unmodified sequences identified with 1% FDR were obtained from (Racle et al., 2019).

### Peptide filtering, allele assignment, and core prediction using the HLA-II ligand predictor

To filter potential contaminants or wrongly identified peptides and to determine allelic restriction and peptide binding cores, the HLA-II ligand predictor MixMHC2pred (v1.2) (Racle et al., 2019) was applied to all phosphorylated peptides for all alleles available in each sample (see Table S3 for HLA typing of each sample). Phosphorylated residues were substituted by the unspecific amino acid ‘X’. To filter potential contaminants or wrongly identified peptides, a %rank cutoff of 5% was applied to the data. Peptides not passing this threshold were not considered in any analysis. The same predictions were performed on random phosphorylated peptides selected from a pool of all known phosphosites of the human proteome (Ullah et al., 2016) with the same length distribution as the phosphorylated HLA-II ligands found by MS. Sequence logos including phosphorylated peptides were drawn with the extended version of ggseqlogo (https://github.com/GfellerLab/ggseqlogo) (Wagih, 2017) and phosphorylated residues are shown in purple (Figure 2). Phosphorylated HLA-II binding motifs shown in Figure 2 were built using only peptides containing phosphorylated residues within the binding core.

### Positional Distribution of phosphorylated residues in HLA-II ligands

The frequency of phosphorylated residues inside of the binding core and in PFRs were compared to the total fraction of residues in these two regions of the peptides, and a two-sided Fisher’s exact test was applied to calculate the P-value (Figure 3A). Only peptides with phosphorylated residues in the first three positions of the N-terminal region or in the last three positions of the C-terminal region were used to compute the distribution of phosphorylated residues in PFRs (Figure 3B). The distribution of phosphorylated residues per position in the core was computed position-wise for all peptides that contained at least one phosphorylation in the binding core (Figure 3C).

### Competition Binding Assays

To test binding of different phosphorylated HLA-II ligands, competition assays were performed for HLA-DRB1*01:01 and HLA-DRB1*07:01 with two and one different peptides detected by MS in the samples, respectively. The competition assays were performed by mixing in v-bottom 96-well plate (Greiner Bio-One) in a citrate saline buffer (100 mM citrate, pH 6.0), with 0.2% β-octyl-glucopyranoside (Calbiochem), 1×complete protease inhibitors (Roche), and 1 μg of the biotinylated empty allele with a FLAG-tagged peptide at fixed concentration of 2 μM (Influenza HA_307-319_ for HLA-DRB1*01:01 and NY-ESO-1_87-99_ for HLA-DRB1*07:01). The peptide of interest was added to this mix into each well at a final concentration of 0, 0.13, 0.41, 1.3, 3.7, 11.1, 33.3, and 100 μM. For the control, untagged peptide (Influenza HA_307-319_ or NY-ESO-1_87-99_) were added at the respective concentrations to the mix of allele and FLAG-tagged peptide. After incubation at 37°C overnight, the binding of the tagged peptides to HLA-II molecule was measured by ELISA. The mix was transferred to a plate coated with avidin and the FLAG-peptide was detected with an anti-FLAG-alkaline phosphatase conjugate (Sigma), developed with pNPP SigmaFAST substrate and absorbance was read with a 405nm – filter (Figure 3D-F).

### Kinase motifs

To detect enrichment of kinase motifs in phosphorylated HLA-II ligands, occurrences of all motifs from the PhosphoMotif Finder of the Human Protein Reference Database (Amanchy et al., 2007) were searched in phosphorylated as well as unmodified HLA-II ligands (Figure 4A). To be able to search each motif on all peptides, including those that had the phosphorylated residue at the first or last positions, each phosphorylated and unmodified peptide was mapped to its source protein and N- and C-terminally extended. To compute frequencies in Figure 4A, occurrences of kinase motifs were normalized by the amount of phosphorylated residues of the corresponding motif in all phosphorylated peptides (e.g., amount of pS and pT in all phosphorylated peptides for motif [D/E][pS/pT]). Similarly, frequencies of kinase motifs in unmodified peptides were determined by normalization with the amount of the unmodified counterpart of the phosphorylated residues of the corresponding motif in all unmodified peptides (e.g., amount of S and T in unmodified peptides for motif [D/E][S/T]). For comparison, the same analysis was also performed on the human phosphoproteome (Sharma et al., 2014). The most common and non-redundant kinase motifs that showed a P-value ≤ 0.05 between phosphorylated and unmodified HLA-II peptides (computed with one-sided Fisher’s exact test) are shown in Figure 4A. To analyze whether the difference in kinase motifs between phosphorylated HLA-II and the human phosphoproteome (Sharma et al., 2014) is due to a gene bias of source proteins, a universal set of source genes of MS-detected sequences was defined. This universal gene set contained all source genes of phosphorylated and unmodified HLA-II sequences, source genes from a phosphoproteome (Sharma et al., 2014) and a MS-based human proteome (Wilhelm et al., 2014). Next, source genes of known phosphosites from the phosphoproteome containing the [pS/pT]P motif were identified and the overlap with unmodified HLA-II ligands was computed (see Figure S2). P-values were computed with one-sided Fisher’s exact tests.

### Predictor

Predictions of interactions between HLA-II alleles and phosphorylated peptides were based on the previously developed HLA-II ligand prediction method MixMHC2pred (Racle et al., 2019). Following our previous work on phosphorylated HLA-I ligands (Solleder et al., 2020), the MixMHC2pred training framework was extended to consider 23 amino acids and the phosphorylated peptides were added to the training set used in (Racle et al., 2019). MixMHC2pred was then retrained on this combined dataset of both phosphorylated and unmodified HLA-II ligands. For the leave-one-sample-out cross-validation, each sample from the dataset was iteratively used as test set and all phosphorylated peptides that were found in this sample were removed from the training data of MixMHC2pred. In each round of the cross-validation, the set of phosphorylated peptides used for training was added to the existing (unmodified) training data previously used (Racle et al., 2019). Five times the amount of positive phosphorylated peptides were added to the testing data as negative peptides. Peptides used as negatives in the test set were of lengths 12 to 25 amino acids and contained a phosphosite from a pool of all known phosphosites of the human proteome (Ullah et al., 2016) (the phosphosite itself, the length of the peptide as well as the position of the phosphosite in the 12 to 25-mer were randomly chosen).

Other existing HLA-II predictors (MARIA (Chen et al., 2019) and NetMHCIIpan4.0 (Reynisson et al., 2020a)) were used to benchmark the prediction results (Figure 4B and 4C). MARIA was used with the unmodified version of the phosphorylated peptides (S, T, Y instead of pS, pT, pY) as well as gene names of the peptides’ source proteins and only applied to HLA-DR alleles. Phosphorylated residues in HLA-II ligands were substituted by ‘X’ for predictions with NetMHCIIpan4.0. For the prediction of peptides coming from samples measured also with pan-HLA-II specific antibodies, alleles available in MixMHC2pred were used for both MixMHC2pred and NetMHCIIpan. For comparison of the predictions with each method, the area under curve (AUC) of the receiver operating characteristic (ROC) was computed for each sample and each predictor.

## Software availability

The command-line script to run the new version the HLA-II ligand prediction method (MixMHC2pred v1.3) is available at https://github.com/GfellerLab/MixMHC2pred/tree/MixMHC2pred1.3.

## Author contribution

D.G designed and supervised the study. M.S. analyzed the data and developed the computational tools. M.B.-S. analyzed the MS spectra. P.G. performed the experiments. J.R. provided support for the data analysis and computational tool developments. G.C. provided technical support. M.S. and D.G. wrote the manuscript and all authors edited the manuscript.

## Declaration of interests

The authors declare no competing interests.

## Notes

### Competing Interest Statement

The authors have declared no competing interest.

