## Supplemental Information for "Deciphering the landscape of phosphorylated HLA-II ligands"

### Supplementary Figures

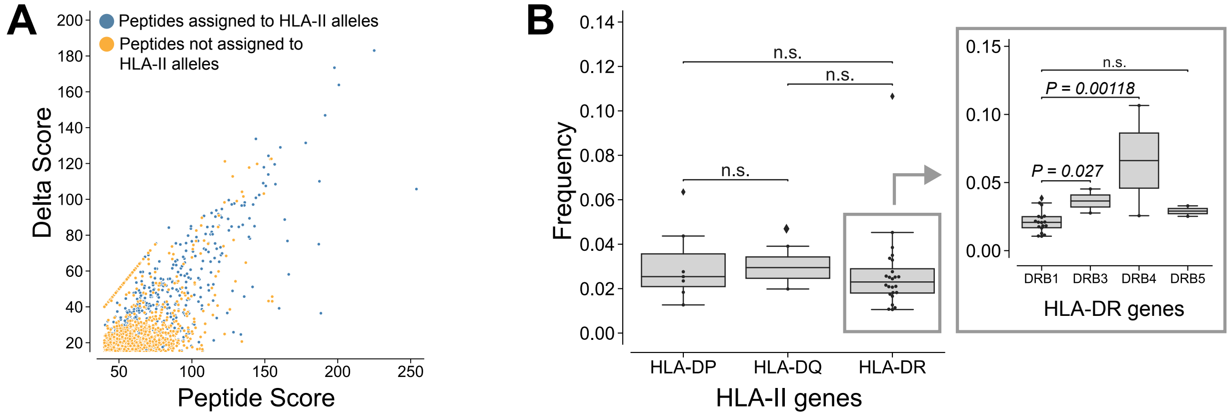


**Figure S1: HLA-II alleles present similar fractions of phosphorylated ligands. (A)** Plot of Andromeda search engine peptide spectrum match scores (*Peptide Score*) vs. score differences to the second-best peptide spectrum match (*Delta Score*) of the phosphorylated peptides. Peptides assigned to HLA-II alleles are shown in blue, the others are shown in orange. **(B)** Frequency of phosphorylated peptides in the HLA-II peptidome for the alleles of each gene (HLA-DRB1/3/4/5, -DP, or -DQ).

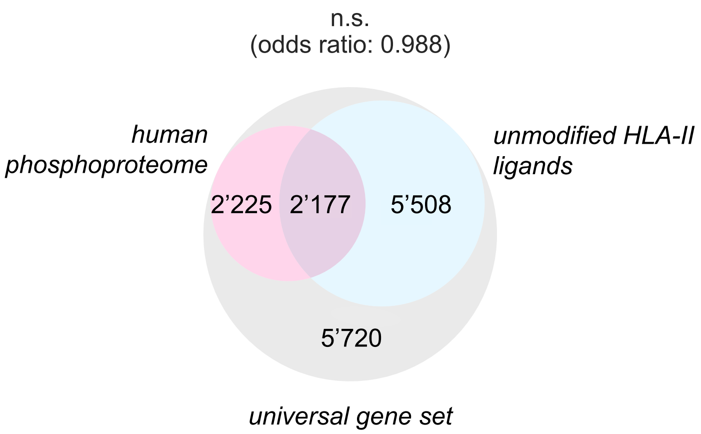


**Figure S2: Overlap between source genes of proteins containing phosphosites with the [pS/pT]P motifs in the human phosphoproteome (pink) and source genes of all HLA-II ligands (light blue).**

### Supplementary Tables

**Table S1: MS/MS data for all phosphorylated peptides used in this study.** The list contains phosphorylated peptides of length 12 to 25, identified across all 23 samples, with a *Peptide Score* ≥ 40, a *Delta Score* ≥ 10, and a localization probability for phosphorylation > 0.75. Phosphorylated residues are given in lowercase letters.

**Table S2: List of phosphorylated peptides that were assigned to HLA-II alleles.** The second column indicates the allele(s) to which the phosphorylated peptide is predicted to bind. The third column shows the predicted core(s). Phosphorylated residues are given in lowercase letters.

**Table S3: List of all samples used in this work.** The second column corresponds the HLA typing of each sample and the third column indicates the HLA alleles available for predictions with MixMHC2pred.
